# Temporal constraints of conscious tactile perception in the primary somatosensory cortex

**DOI:** 10.1101/2025.08.21.671533

**Authors:** Louisa Gwynne, Luigi Tamè

**Author notes:** Address for correspondence: Louisa Gwynne or Luigi Tamè School of Psychology University of Kent, CT2 7NP, Canterbury, United Kingdom.

## Abstract

The primary somatosensory cortex (S1) has long been implicated in tactile perception, yet its precise role in conscious tactile detection remains uncertain. The current study investigated the causal and time-specific involvement of S1 in tactile detection using single-pulse transcranial magnetic stimulation (spTMS). In two experiments, spTMS was applied over contralateral S1, an active control site (inferior parietal lobe; IPL), or under a sham condition at short (25 & 75 ms; Experiment 1) and longer (130 ms; Experiment 2) intervals following electrotactile stimulation of the finger. Participants performed a go/no-go detection task at sensory threshold. In Experiment 1, tactile sensitivity was significantly reduced following early S1 stimulation compared to both active control and sham conditions. However, no such effect was observed in Experiment 2, indicating a temporally limited role of S1 in conscious detection. Moreover, self-reported TMS-related distraction ratings did not account for the observed sensitivity differences, suggesting sensitivity-specific modulation by early TMS rather than general task disruption. These findings support a causal role for early S1 activity in conscious tactile detection. We propose that disruption at this early stage interferes with the initial encoding of tactile input, thereby attenuating not only immediate perceptual awareness, but also subsequent functions such as discrimination and retention. Overall, the results underscore the constrained role of S1 in conscious stimulus detection and highlight the importance of neural networks beyond S1.

## Introduction

The ability to detect touch supports our fundamental need to interact with the external world. Early primate studies showed that primary somatosensory cortex (S1) receives tactile input via thalamocortical projections to Broadmann areas (BA) 3b and 1 (Lamotte & Mountcastle, 1979). These signals arrive approximately 20 milliseconds (ms) after contralateral median nerve stimulation and around 20-25 ms after digit nerve stimulation, marking the earliest cortical processing of tactile input (Allison et al., 1991; Peterson et al., 1995; Tsujinaka et al., 2023). Moreover, somatosensory-evoked responses in S1 have been associated with perceptual detection (e.g. Jones et al., 2007; Palva et al., 2005). Nonetheless, the precise role of S1, encompassing the specifics of what it does and how it operates remains open and disputed.

The earliest component of the somatosensory evoked potential (SEP) following peripheral stimulation is reflected in the N20 component in electroencephalography (EEG) or the M20 component in magnetoencephalography (MEG). This early negative potential is generated in BA 3b, contralateral to the locus of stimulation (Allison et al., 1991; Mauguière et al., 1997). Although the N20/M20 is widely considered the first cortical response to tactile input (Allison et al., 1991; Desmedt & Tomberg, 1989; Peterson et al., 1995), its role in conscious detection remains debated. Seminal work by Libet and colleagues demonstrated that such early cortical responses do not reflect conscious awareness of the stimulus (Libet et al., 1967). In contrast, the slightly later P50 component – primarily originating from BAs 1 and 3b (Allison et al., 1992) – has been found to correlate with tactile detection (Soininen & Järvilehto, 1983). Furthermore, M/EEG components peaking between 30 and 70 ms post-stimulus have been shown to predict tactile detection performance in both non-human primates (Cauller & Kulics, 1991) and humans (Hirvonen & Palva, 2016; Jones et al., 2007; Palva et al., 2005).

In contrast to the view that early components originating in S1 reflect stimulus detection, other research suggests these responses instead represent pre-conscious sensory processing rather than perceptual awareness (Schubert et al., 2006; Wühle et al., 2010, 2011). In such view, later SEP components occurring after 100 ms, such as the P100 and N140, are more consistently associated with conscious detection, reliably distinguishing between detected and undetected target (Auksztulewicz et al., 2012; Forschack et al., 2020; Schröder et al., 2021; Schubert et al., 2006; Wühle et al., 2011). Both components have origins in the secondary somatosensory cortex (S2), with the P100 also originating in S1 (Allison et al., 1992; Mauguière et al., 1997). Taken together, this evidence highlights ongoing uncertainty regarding S1’s contribution to detection-particularly the latency at which the tactile signal is key for conscious perception.

In addition to M/EEG research, several transcranial magnetic stimulation (TMS) studies have investigated the causal role of S1 in tactile detection (Cohen et al., 1991; Hannula et al., 2005; Harris et al., 2002; Seyal et al., 1997; Tamè & Holmes, 2016). A seminal study by Cohen and colleagues (1991) demonstrated that single-pulse TMS (spTMS) over the contralateral sensorimotor cortex supressed electrotactile detection 200ms before and 20 ms after stimulus onset (Cohen et al., 1991). Similarly, spTMS over contralateral S1 was found to impair the detection of electrotactile stimulus trains when administered 100 ms before and 20 ms after touch (McKay et al., 2003), suggesting a contribution of early S1 processing to tactile detection. However, later studies yielded inconsistent findings. For instance, spTMS over S1 failed to attenuate tactile detection, a result attributed to the absence of simultaneous motor cortex stimulation present in earlier studies (Koch et al., 2006). Nonetheless, magnetic resonance imaging (MRI)-navigated TMS applied to S1 at 20 or 50 ms post-stimulus was shown to impair detection (Hannula et al., 2005), reinforcing the potential contribution of S1 involvement in conscious tactile detection.

In a recent TMS study, mechanical tactile detection was reportedly impaired by two successive sp-TMS pulses delivered over contralateral S1 25 and 75 ms after stimulus onset (Tamè & Holmes, 2016), relative to TMS over an active control site. Notably, this effect was present only in a one-interval ‘yes/no’ design task, but not in a two-interval forced choice paradigm requiring the identification of a target in the first or second interval. This finding suggests that the contribution of S1 to tactile detection may depend on the cognitive demands of the task (Tamè & Holmes, 2016; see, also, Katus et al., 2015). Moreover, the temporal window of S1 involvement in conscious detection may differ as a function of the stimulation type, namely mechanical or electrical (Alouit et al., 2025). What is more, the temporal dynamics of its contribution to conscious detection remain largely unexplored.

Beyond detection, it is well-evidenced that S1, in both monkeys and humans, contributes to later tactile processes of higher-order phenomena including, such as discrimination (Hannula et al., 2008; Harris et al., 2002; Kulics, 1982; Kulics & Cauller, 1986; Lamotte & Mountcastle, 1979; Morley et al., 2007; Mountcastle et al., 1969; Romo et al., 2000), as well as, interhemispheric integration (Tamè, Pavani, Papadelis, et al., 2015), retention (Harris et al., 2001, 2002; Katus et al., 2015), sensorimotor integration (Tamburin et al., 2001; Tamè, Pavani, Braun, et al., 2015) and localisation (Braun et al., 2005; Harris et al., 2001; Iwamura et al., 2002; Katus et al., 2015). Moreover, M/EEG studies tend to associate detection with later processing (beyond the N20), however, TMS evidence points to an earlier, causal role. This underscores the uncertainty surrounding the timing of S1’s involvement in tactile detection, and most importantly, its key contribution to tactile awareness.

In the present study, we employed a spTMS paradigm in which TMS was delivered to contralateral S1, 25 and 75 ms (Experiment 1) or, 130 ms (Experiment 2) after the onset of electrotactile stimulation at sensory threshold to the index finger during a go/no-go task. The effect of spTMS over S1 was compared to an active control site (inferior parietal lobe) and a sham condition, in which the coil was flipped over the vertex. It was hypothesised that causal involvement of S1 to stimulus detection would be marked by significantly reduced stimulus sensitivity compared to both the active control and sham TMS conditions (Figure 1). In-line with both TMS and M/EEG literature, we predicted significantly reduced tactile sensitivity in both Experiment 1 and 2 by TMS over S1.

**Figure 1.**
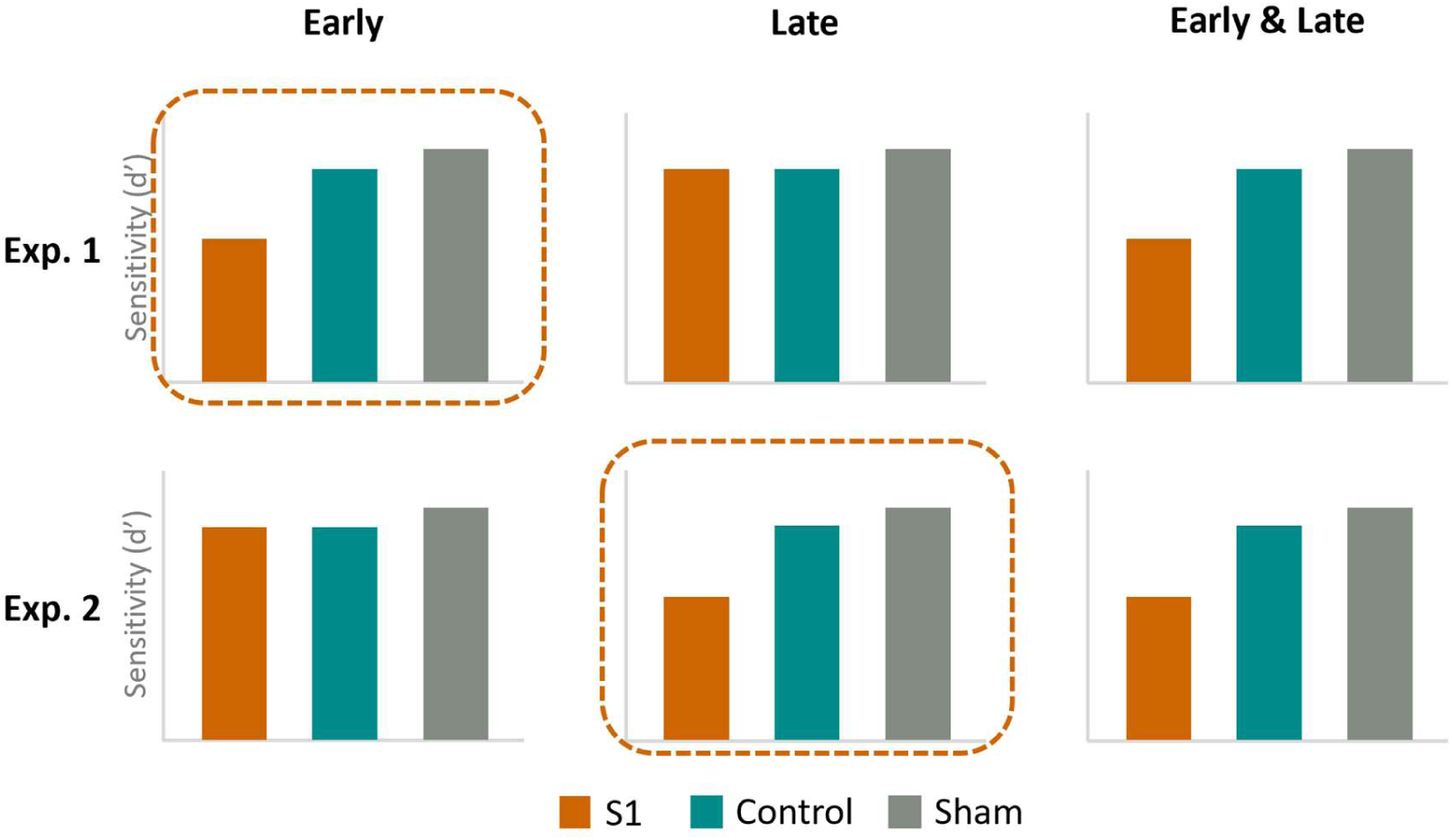
Involvement of S1 in conscious tactile detection. Examples of the predicted tactile sensitivity (d’) as a function of spTMS stimulation conditions: S1, IPL, or, vertex (sham) for Experiment 1 (early TMS <100 ms) and 2 (late TMS: > 100 ms). Dashed orange boxes represent predicted results for Experiment 1 and 2, showing significant involvement of S1 to tactile detection by reduced sensitivity (d’) compared to control and sham TMS at the early and later stages of tactile representation processing.

## General method and materials

### Participants

An a priori power analysis using G*Power (Faul et al., 2007) estimated that a sample of 19 participants would be sufficient to achieve 80% (alpha = 0.05) to detect a medium-to-large effect (dz = 0.6) in a within-subjects t-test comparing tactile sensitivity (d’) between the experimental and control TMS sites. To mitigate for possible dropout and adequate performance at the task we aimed for a sample of 25 participants. In Experiment 1, 3 participants were excluded from analysis due to below chance performance in at least one of the TMS conditions, leaving a final sample of twenty-two participants (Mean_Age_= 20.23, SD_Age_= 5.28 years; 4 males, 18 females). Twenty participants were right-handed and 2 left-handed as indicated by self-report on the Edinburgh Handedness Inventory questionnaire (EHI; Oldfield, 1971; M = 72.4, range = –55.56 – +100). In Experiment 2, twenty-two new participants were tested matching the sample size of Experiment 1, however, 2 were removed for below chance performance in at least one of the TMS conditions, leaving a sample of twenty (Mean_Age_= 22.9. SD_Age_= 7.55 years; 5 males, 15 females). Nineteen were right-handed and 1 left-handed (EHI; M= 81.54, range = –75 – +100). All participants were recruited voluntarily in exchange for monetary payment at a rate of £7.50 an hour or via a university participation scheme for course credit. Participants had normal sensation of the hands and arms as indicated through free self-report. Participants were screened for TMS contraindications, with all reporting no neurological or psychiatric conditions, and no current use of psychiatric and neuroactive medications. All procedures were approved by the departmental ethics committee at the University of Kent and adhered to published TMS safety guidelines (Rossi et al., 2021).

### Tactile stimuli

Tactile stimulation consisted of a single transcutaneous electrical stimulation delivered through two stainless steel electrode rings, fitted at the proximal and intermediate phalange of the participants’ right index finger, connected to a bipolar constant current stimulator (DS5; Digitimer, Welwyn Gardne City, United Kingdom), controlled using a custom-made programme in MATLAB (version 2019b) through a National Instruments data-acquisition card (6001). Square wave pulse width was set a 0.2 ms. At low intensities (<∼6 mA) this created a singular, fast onset-offset tap-like sensation by momentary activation of predominantly Aβ mechanoreceptive fibres. The right hand was always covered from site throughout the experiment to avoid uncontrolled effects caused by the vision of the body (Cardini et al., 2011; Tamè et al., 2017).

### Tactile threshold

Tactile stimuli were delivered at the participants’ electrotactile detection threshold, characterised as the minimum stimulator output detectable (Table 1). This was estimated with an automated 2-down/1-up staircase procedure converging to an approximately 71% perceivability (Levitt, 1971). Starting at 0.1 mA, the stimulation increased by 0.3 mA step-sizes, reduced to 0.1 mA after the first reversal and to 0.02 mA after the second reversal. The staircase ended at the eighth reversal and the sensory detection threshold (SDT) was calculated as the average of the last two reversals. This procedure was determined by a series of pre-experiment pilots showing that this produced a stimulus 70-80% perceivable on 10 out of 20 trials. Electrotactile detection thresholds did not differ between participants in the two experiments (*t* = 0.001, *p* = .999).

**Table 1.** Resting motor thresholds (RMT) and sensory detection thresholds (SDT) across experiments.

| Participant | RMT (% MSO) |  | SDT (mA) |  |
| --- | --- | --- | --- | --- |
|  | Exp 1. | Exp 2. | Exp 1. | Exp 2. |
| 1 | 40.5 | 43.0 | 1.53 | 1.68 |
| 2 | 34.0 | 49.5 | 0.83 | 0.95 |
| 3 | 47.5 | 35.0 | 1.83 | 1.45 |
| 4 | 38.0 | 48.0 | 1.79 | 1.23 |
| 5 | 40.5 | 47.5 | 1.82 | 1.53 |
| 6 | 37.5 | 44.5 | 1.63 | 2.34 |
| 7 | 40.5 | 41.5 | 2.26 | 1.63 |
| 8 | 42.5 | 36.0 | 0.94 | 1.48 |
| 9 | 41.5 | 39.0 | 2.03 | 1.19 |
| 10 | 48.5 | 51.5 | 1.30 | 1.25 |
| 11 | 37.5 | 37.5 | 1.21 | 1.75 |
| 12 | 42.0 | 46.0 | 1.61 | 2.36 |
| 13 | 33.0 | 41.5 | 1.55 | 2.19 |
| 14 | 39.5 | 41.0 | 1.46 | 2.44 |
| 15 | 45.5 | 47.5 | 1.45 | 1.65 |
| 16 | 41.5 | 45.0 | 1.24 | 1.33 |
| 17 | 47.5 | 38.5 | 2.50 | 0.75 |
| 18 | 38.5 | 40.5 | 2.17 | 0.96 |
| 19 | 50.0 | 55.0 | 1.56 | 1.53 |
| 20 | 42.0 | 41.5 | 2.10 | 1.69 |
| 21 | 43.5 | - | 1.23 | - |
| 22 | 44.5 | - | 1.19 | - |
| Mean | 41.64 | 43.48 | 1.6 | 1.57 |
| SD | (4.43) | (5.29) | (0.43) | (0.48) |

**Note. MSO = Maximum stimulator output (%)**

### TMS and Neuronavigation

Biphasic single-pulse TMS (spTMS) was delivered with a MAG and MORE PowerMAG 100 Repetitive Magnetic Stimulator via a 100 mm diameter figure-of-eight coil. Electromyography (EMG) was recorded with two Ag-AgCI surface electrodes placed over the first dorsal interosseus (FDI) muscle of the right hand in a belly-tendon montage with a ∼2cm inter-electrode distance. The ground electrode was placed 1-2 cm proximal to the right pisiform bone of the left wrist and skin preparation was completed using an alcohol swab. EMG signal was sampled in BrainSight Neuronavigation software (version 2.4.10; Rogue Research Inc., Montreal, QC) with a 3000 Hz sampling frequency. Scalp localisation was optimised using the same BrainSight Neuronavigation software. The total number of TMS pulses delivered to each participant was between 220-480 (including M1 localisation and TMS threshold procedures).

### M1 localisation and RMT

TMS intensity was delivered at 120% of the resting motor threshold (RMT), defined as the optimal scalp positioning to elicit at least a 50-mV peak-to-peak motor-evoked potential (MEP) in the right FDI muscle while at rest in five out of 10 trials (Rossini et al., 1994). A 4×4 square grid with 10 mm spacing was drawn onto the participant’s reconfigured digitalised scalp in BrainSight, centred on our lab average left M1-FDI coordinates from previous experiments (Gwynne & Tamè, 2025, bioRxiv). TMS was applied at 35% of the maximum stimulator output, increasing in increments of 5% until muscle evoked activity was visible on the EMG and the coil was moved across the grid to find the grid point with the maximum MEP output. TMS intensity was then adjusted until desired MEPs were recorded. The coil handle was positioned 45 degrees relative to the sagittal plane. Across both experiments, the mean average right M1-FDI RMT was 42.51 % (SD = 4.89; Table 1) and the mean average right M1-FDI MNI coordinates were –37.76, –9.29, 59.06 (SD= 3.65, 6.35, 3.02; x, y, z).

### S1 Localisation

Left S1 was localised relative to the left M1-FDI hotspot, estimated in BrainSight Neuronavigation by moving the left M1-FDI scalp coordinates 20 mm along the X axis and –5 mm along the Y axis. This protocol was made in consideration of a systematic review finding that S1 lies approximately 2 cm lateral and 0.5 cm posterior to the FDI motor hotspot (Holmes et al., 2019; Holmes & Tamè, 2019). The coil handle was held at a 90-degree angle relative to the sagittal plane as this has been shown to be an effective orientation to maximise S1 stimulation and minimise unwanted M1 responses (Tamè & Holmes, 2023). Across both experiments the average S1 MNI coordinates were –46.47, –16.36, 54.1 (SD= 2.58, 5.6, 2.82; x, y, z; Figure 2).

**Figure 2.**
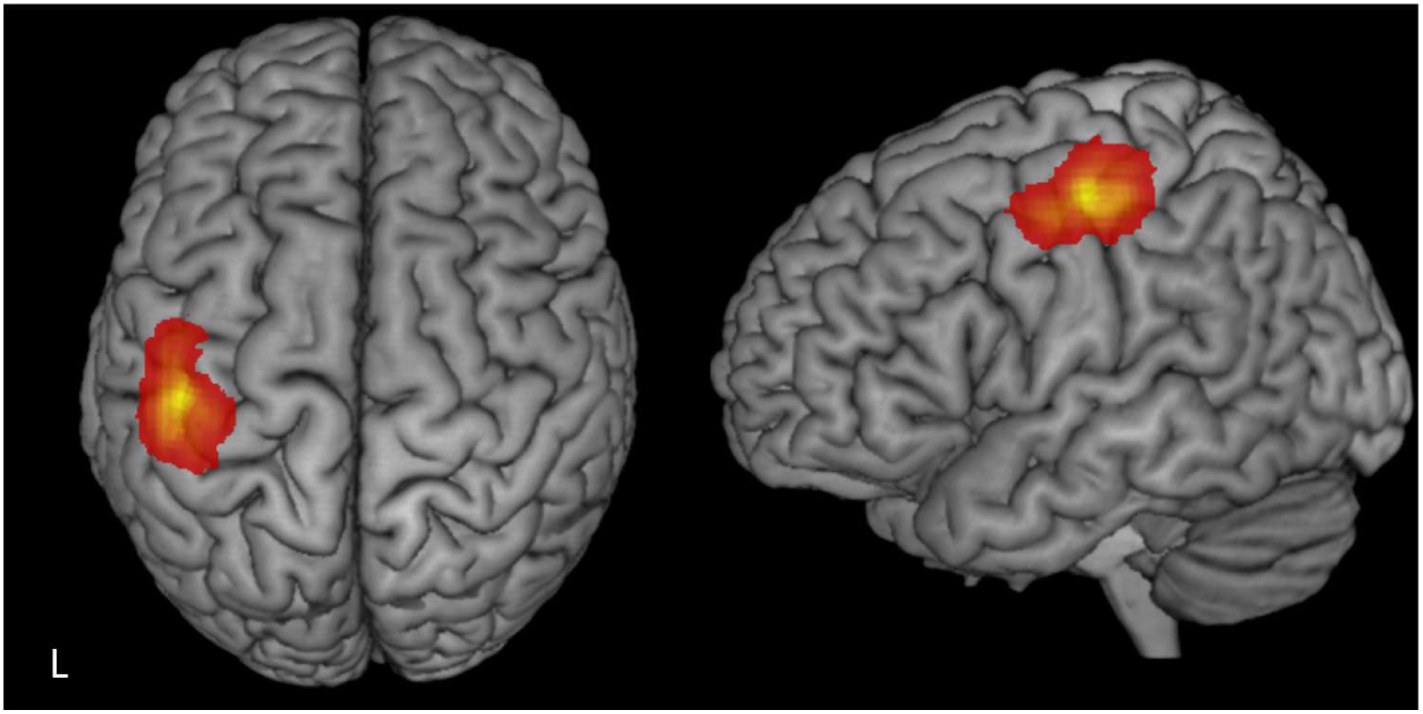
Showing determined left S1 MNI coordinates across participants and experiments. Heatmaps were made using custom MATLAB scripts and SPM12. Participants’ MNI coordinates were transformed into voxel space of an anatomical template. A blank 3D volume with the same dimensions as the template was then initialized and each voxel correspondigng to a coordinate was incremeted by one to reflect any frequency of overlap of coordinates. Lastly the density image was smoohed by a 6mm FWHM guassian kernal.

### Sham and control sites

In sham conditions, the coil was held over the vertex with a handle position 180 degrees relative to the sagittal plane and the coil was flipped with the active side facing upwards so that the magnetic field was dissipated into the air (Figure 3C). The left inferior parietal lobe (IPL) acted as the active control site using the average MNI co-ordinates (–56, –56.1, 36.2) from Tamè and Holmes (2016). Whilst stimulation of this site produces TMS-induced artefacts comparable to S1, a previous study showed IPL did not elicit any significant fMRI activation during tactile detection (Tamè & Holmes, 2016). The coil handle was positioned at a 90-degree angle relative to the sagittal plane. However, coil orientation was adjusted at the start of the main experimental task to match the level freely reported TMS-related discomfort between S1 and IPL. On average, this resulted in a slight forward tilt of the coil, lifting the lower edge away from the scalp to minimize discomfort induced by facial or scalp muscle contractions.

**Figure 3.**
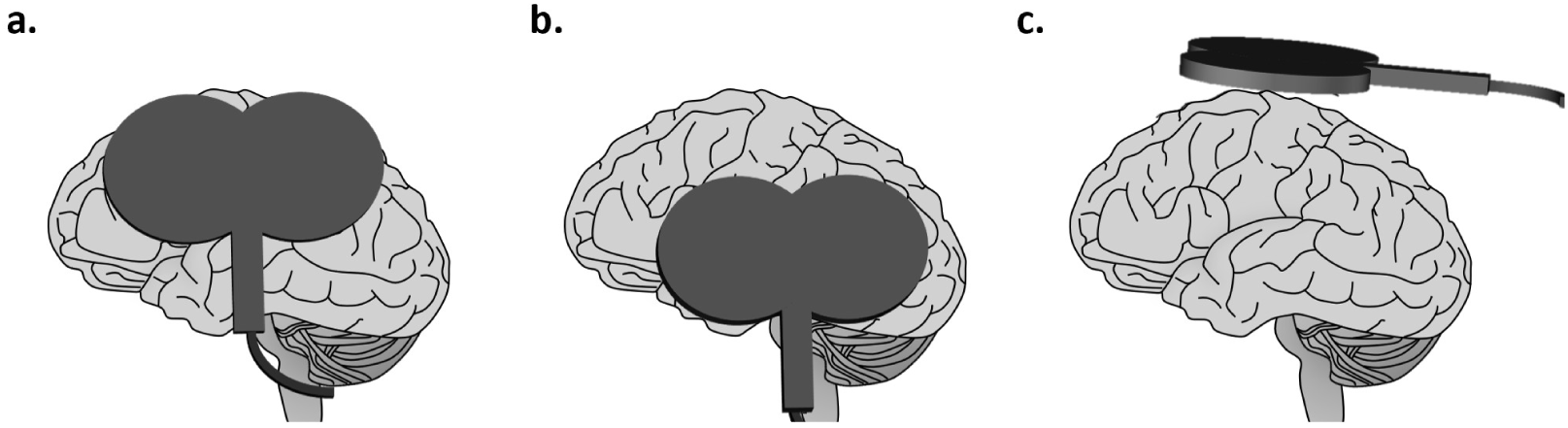
TMS conditions and coil location and orientation over scalp. a) Experimental target primary somatosensory cortex (S1), b) active control Inferior parietal lobe (IPL) and sham vertex with a flipped coil directed away from the scalp.

## General procedure

### Experimental Task

After completing TMS thresholding and localisation, followed by the tactile thresholding procedure, all participants completed a tactile detection Go/No-go task. Participants sat with their right hand and forearm resting on a desk and fixed their gaze on a white fixation cross centrally positioned on a monitor 60 cm from the participant. At the start of a trial, the fixation cross turned black for four seconds, after which the cross turned back to white to indicate the end of a trial. The inter-trial interval (ITI) was 2.5 seconds in order to prevent any carryover effects of tactile stimulation or spTMS from the previous trial. The task consisted of six blocks of 20 trials. A tactile stimulus was pseudorandomly presented on fifty per cent of trials (10/20); a 0mA stimulus intensity was delivered on trials with no tactile stimulus. Tactile stimulus presentation occurred at a randomly jittered timepoint between 1-2 seconds from the start of the trial. Participants were instructed that on every trial they may or may not feel a stimulation on the index finger and to press the “0” key with the left index finger when they felt a stimulus. The left index finger was rested on the “0” key throughout. No response was required if no stimulus was perceived. spTMS was delivered on every trial at 120% of the M1-FDI RMT. The Timepoint of TMS stimulation varied across experiments (Figure 4). The location of TMS (S1, control or sham) was counterbalanced across blocks under an ABCCBA counterbalanced sequence to which participants were randomly assigned. 2-minute breaks were taken between experimental blocks to allow coil re-positioning and participant rest (total time to complete the task was about 25 minutes). No feedback on accuracy was provided.

**Figure 4.**
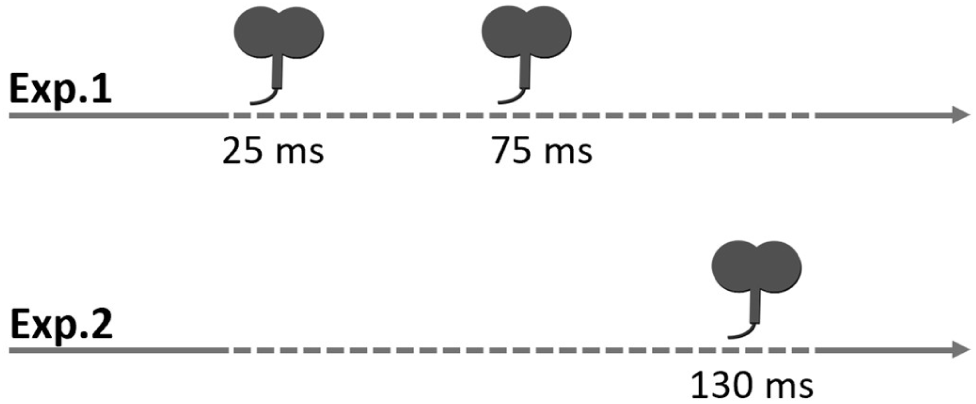
Schematic depiction of TMS pulse(s) timing relative to tactile onset in Experiments 1 and 2.

## Experiment 1

Experiment 1 investigated the effect of double spTMS over contralateral S1 on tactile detection compared to sham and control TMS. spTMS was delivered 25 and 75 ms after tactile onset. Two pulses were chosen to improve precision of targeting the tactile trace arising in S1 following afferent delivery given that signal arrival time by median nerve stimulation is thought to be around 20 ms (Allison et al., 1991) but, has also been associated with perceptual detection around 70 ms after touch (Palva et al., 2005). Furthermore, note that the effect of single or double pulse TMS has not been found to differently affect tactile detection performance (Tamè & Holmes, 2016).

## Results

Hits were defined as a “yes” response on target-present trials, while false alarms were defined as a “yes” response on target-absent trials. Sensitivity was measured using d-prime, calculated as *d’* = *z*(Hits) – *z*(False Alarms). Response bias was measured using the criterion, calculated as *c* = –0.5 x [*z*(Hits) + *z*(False Alarms)]. To enable Z-score transformations, a continuity correction was applied such that hit and false alarm rates of 1 and 0 were adjusted to 0.99 and 0.1, respectively.

As shown in Figure 5 (left panel), one-tailed paired sample t-tests revealed tactile sensitivity (d’) was significantly reduced when TMS was delivered to S1 (M ± SE d′ = 2.14 ± 0.21) compared to IPL (M ± SE d′ = 2.6 ± 0.29; *t*(21) = –1.88, *p* = .037, *dz* = –0.38) and sham TMS (M ± SE d′ = 2.62 ± 0.21; *t*(21) = –2.67, *p* = .007, *dz* = –0.48). Differently, sensitivity was comparable between IPL and sham TMS (*t*(21) = –0.09, *p* = .931, *dz* = –0.02). Criterion did not significantly differ when TMS was delivered to S1 (M ± SE c = 0.15 ± 0.11) compared to IPL (M ± SE c = 0.25 ± 0.11; *t*(21) = –0.88, *p* = .391, *dz* = –0.2. However criterion at S1 just reached significant difference when compared to sham TMS (M ± SE c = 0.37 ± 0.13; *t*(21) = –2.07, *p* = .051, *dz* = –0.41) and, criterion was significantly lower at IPL than sham TMS (*t*(21) = –2.12, *p* =.046, *dz* = –0.21; Figure 5).

**Figure 5.**
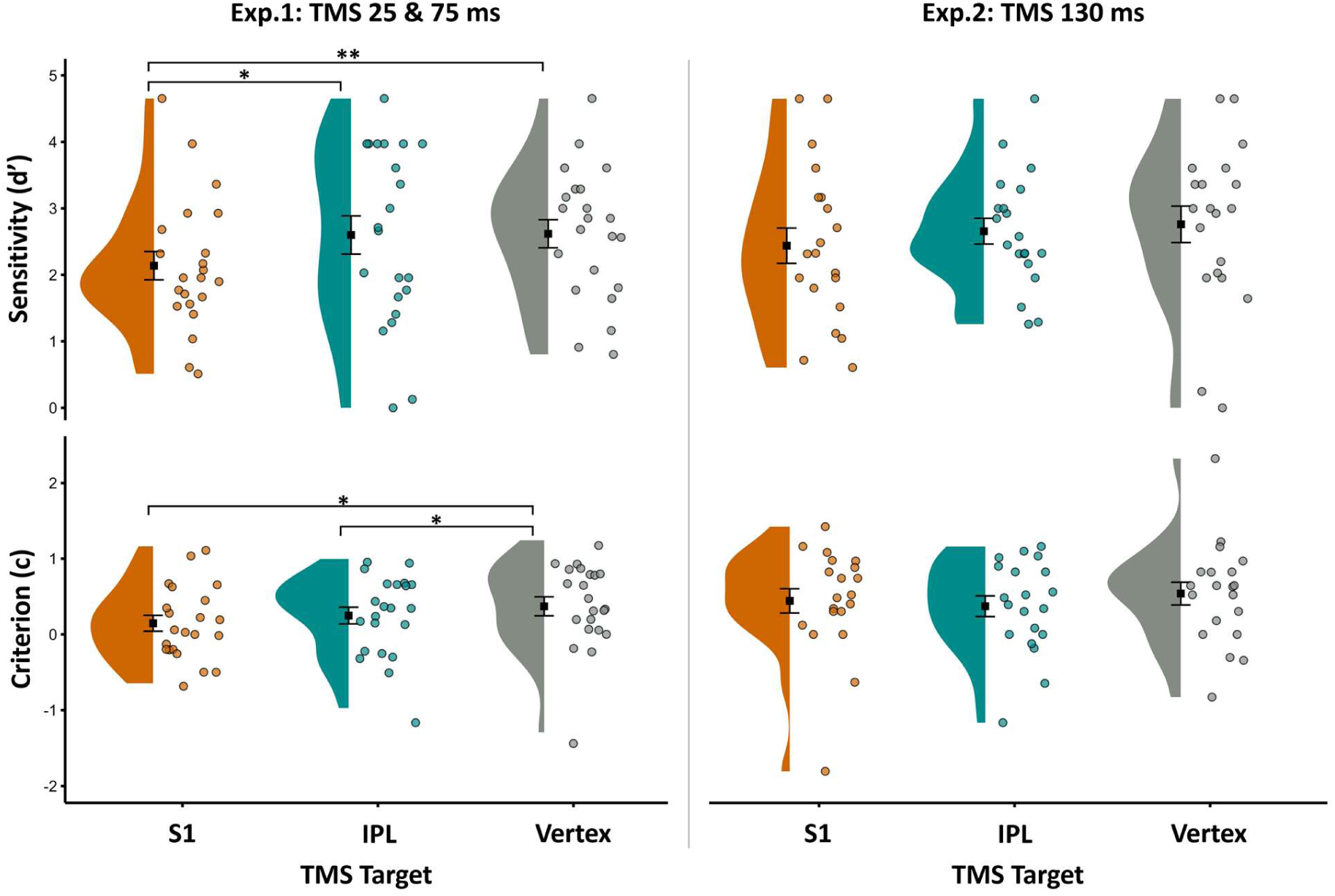
Sensitivity (d’) and criterion (c) as a function of TMS target when early TMS was delivered after afferent onset (25 & 75 ms, Experiment 1) or, late TMS delivery (130 ms, Experiment 2). Black squares show mean and standard error means (SEM). Half violins show data distribution, coloured circles are jittered individual participant data points\**P* ≤ .05, \*\**P* ≤ .01, \*\*\**P* ≤ .001.

Self-reported TMS distraction for each TMS site was averaged across the two blocks of stimulation. Distraction was significantly higher at S1 (M ± SE = 4.76 ± 0.35) than at IPL (M ± SE = 4.22 ± 0.34; *t*(21) = 2.74, *p* =.012, *dz* = 0.34). Differently, distraction at vertex (M ± SE d′ = 2.66 ± 0.23) was significantly lower than both S1 (*t*(21) = 7.31, *p* = <.001, d = 1.44) and IPL (*t*(21) = 6.14, *p* = <.001, *dz* = 1.07. Reaction time (RT) for “hits” on stimulus present trials (“go-trials”) were averaged across the two blocks of stimulation for each TMS site. RT did not significantly differ between S1 (M ± SE RT = 808.03 ± 47.64 ms) and IPL (M ± SE RT = 802.95 ± 45.87 ms; *t*(21) = 0.13, *p* =.896, *dz* = 0.02). However, RTs at vertex (M ± SE RT = 731.71 ± 46.02 ms) were significantly lower than at S1 (*t*(21) = 2.55, *p* = .019, d = 0.35) and at IPL (*t*(21) = 2.18, *p* = .040, *dz* = 0.33). Distraction and reaction time graphs are in Supplementary Materials (Figure S1).

## Experiment 2

Experiment 1 investigated the effect of early spTMS over contralateral S1 on tactile detection compared to sham and control TMS. However, the timeline of this contribution is unclear. Previous studies show that S1 activity persists at least 60 ms after tactile offset (Allison et al., 1992; Mauguière et al., 1997) and, its signal recovery time has been reported to be around 110 ms (Hamada et al., 2002). Moreover, late (>100 ms) activity in the somatosensory cortices has been associated with tactile detection performance (Schubert et al., 2006). Therefore, to investigate any ongoing contribution of S1 to perceptual detection beyond the tested critical window (< 100 ms), in Experiment 2, spTMS was delivered 130 ms after tactile onset.

## Results

As shown in Figure 5 (right panel), one-tailed paired sample t-tests revealed that tactile sensitivity did not significantly differ when TMS was delivered over S1 (M ± SE d′ = 2.44 ± 0.27) compared to IPL (M ± SE d′ = 2.66 ± 0.19; *t*(19) = –0.98, *p* = .169, *dz* = –0.2) and, when compared to sham TMS (M ± SE d′ = 2.76 ± 0.28; *t*(19) = –1.16, *p* = .131, *dz* = –0.27). Sensitivity was also comparable between the active control and sham sites; IPL and sham TMS (*t*(19) = – 0.46, *p* = .648, *dz* = –0.09). Criterion did not significantly differ when TMS was delivered over S1 (M ± SE c = 0.44 ± 0.16) compared to IPL (M ± SE c = 0.37 ± 0.14; *t*(19) = 0.76, *p* = .459, *dz* = 0.01) nor, sham TMS (M ± SE c = 0.54 ± 0.15; *t*(19) = –0.74, *p* = .471, *dz* = –0.14). Furthermore, criterion did not differ between IPL and sham TMS (*t*(19) = –1.68, *p* = .110, *dz* = –0.26; Figure 5).

Self-reported TMS distraction for each TMS site was averaged across the two blocks of stimulation, as in Experiment 1. Distraction did not significantly differ between S1 (M ± SE = 3.78 ± 0.29) and IPL (M ± SE = 3.53 ± 0.37; *t*(19) = 0.89, *p* = .387, *dz* = 0.16). However, distraction was significantly lower for sham TMS (M ± SE = 2.2 ± 0.27) compared to TMS over either S1 (*t*(19) = 7.53, *p* = <.001, *dz* = 1.25) and IPL (*t*(19) = 3.87, *p* = <.001, *dz* = 0.90. Lastly, RT did not significantly differ between S1 (M ± SE RT = 714.57 ± 52.71 ms) and IPL (M ± SE RT = 689.07 ± 40.59 ms; *t*(19) = 0.70, *p* =.493, *dz* = 0.12) nor S1 and sham TMS (M ± SE RT = 642.39 ± 55.36 ms; *t*(19) = 1.79, *p* = .089, *d* = 0.3) and, did not differ between IPL and sham (*t*(19) = 1.49, *p* = .153, *dz* = 0.19).

## Discussion

This study investigated the temporal dynamics of primary somatosensory cortex involvement in tactile detection using inhibitory single-pulse transcranial magnetic stimulation. Tactile sensitivity was significantly reduced when TMS was applied over the contralateral S1 shortly after tactile stimulus onset (25 & 75 ms; Experiment 1), but not when applied at a later time (130 ms; Experiment 2). This effect was specific to S1, as no comparable reduction in sensitivity was observed when spTMS was applied over an active control site (IPL) or using sham TMS (over vertex with a flipped coil). These findings strongly support the notion that S1 plays a critical, yet temporally constrained, role in conscious tactile stimulus detection reflecting its key role in the initial encoding.

Our findings of early but not late S1 involvement in tactile detection stand somewhat in contrast to M/EEG research, which has demonstrated late S1 activity is associated with conscious detection. Specifically, the P100 and N140 components have been shown to correlate with conscious stimulus detection (Auksztulewicz et al., 2012; Forschack et al., 2020; Schröder et al., 2021; Schubert et al., 2006; Wühle et al., 2011), suggesting that late S1 activity contributes to tactile detection, at least to some extent. To reconcile these findings with the present results, we propose that the early causal contribution of S1 reflects its role in the initial encoding of the tactile signal. This in turn provides a neural representation of touch that can immediately or later be distributed across cortical networks to support perceptual functions such as tactile detection, discrimination, retention and learning. Indeed, disruption of the signal during this early time window may impair both immediate and later processes related to perceptual and cognitive functions that requires the tactile sensory signal. Supporting this view, previous studies have shown that TMS over contralateral S1 disrupts performance in tasks involving vibrotactile frequency discrimination and tactile working memory (Harris et al., 2002; Morley et al., 2007), as well as temporal discrimination (Hannula et al., 2008). Moreover, the temporal specificity of S1 involvement observed in our study likely reflects a broader neural architecture underlying tactile processing that extends beyond S1. This interpretation is consistent with the findings of Tamè and Holmes (2016), who demonstrated that the contribution of S1 to tactile detection is shaped by task demands.

The observed early involvement of S1 in tactile detection aligns with previous studies showing that TMS over the contralateral somatosensory cortices impairs detection when applied 20-50 ms after electrotactile stimulation (Cohen et al., 1991; Hannula et al., 2005; McKay et al., 2003; Seyal et al., 1992). However, unlike earlier work, the present study controlled for non-specific TMS effects such as auditory and somatosensory artefacts, by including an active control site, consistent with recent methods used by Tamè and Holmes (2016). Given that higher TMS intensity can increase tactile suppression (McKay et al., 2003), a finding possibly attributable to non-specific TMS effects, the current study also collected subjective TMS-related distraction ratings. In Experiment 1, expectedly, S1 and IPL were rated as more distracting than sham TMS. However, unexpectedly, S1 was rated as significantly more distracting than IPL. Despite this, the reduction in tactile sensitivity following S1 TMS cannot be attributed solely to distraction. If distraction inherently impaired tactile sensitivity, IPL should also have led to reduced sensitivity compared to sham, given that it was also rated as more distracting than sham. Instead, sensitivity was comparable between IPL and sham conditions. We propose that the higher distraction ratings for S1 may reflect a post hoc attribution of task difficulty, where participants’ distraction ratings were influenced by perceived decreased performance. This interpretation is further supported by Experiment 2, in which no differences in either sensitivity or distraction ratings were observed between S1 and IPL.

Our findings on the key role of S1 in tactile sensitivity align with those of Tamè and Holmes (2016), who reported reduced vibrotactile sensitivity when TMS was applied over S1 25 and 75 ms post-stimulus onset during a tactile yes/no task. However, differently to our results, which showed that S1 and IPL TMS led to a lower response criterion compared to sham, Tamè and Holmes (2016) found a higher criterion (more conservative responding), following S1 and IPL. We note, however, that this discrepancy may be partly explained by methodological differences. Their study used a no-TMS condition with the coil held away from the head; contrastingly, we administered sham TMS with a rotted coil held on the scalp over the vertex.

The absence of significant differences in criterion across TMS sites in Experiment 2 may be explained by the longer inter-stimulus interval (130 ms), occurring outside of the critical window for tactile detection. At this later stage, perceptual decisions may already be committed, making participants less susceptible to shifts in response strategy. This further supports the idea that S1 involvement in tactile detection is temporally constrained.

Overall, our results demonstrate that tactile detection arises from temporally staged involvement of S1. Importantly, we demonstrate that the effect of S1 TMS is specific to perceptual sensitivity rather than due to non-specific TMS effects. Within the context of the current methodology, this supports a causal role for S1 in tactile detection, which we propose arises from disrupted initial encoding. This disruption likely reduces the probability that the stimulus reaches higher level stages as well as perceptual awareness.

## Credit author statement

Louisa Gwynne: Conceptualization, Methodology, Validation, Formal analysis, Investigation, Writing – original draft, Visualization; Luigi Tamè: Conceptualization, Methodology, Validation, Formal analysis, Resources, Writing, review & editing.

## Acknowledgments

We would like to thank Nicholas Paul Holmes for his insightful feedback on the data and John Allen for his technical assistance with programming the TMS. LG was supported by the South East Network for Social Sciences (SeNSS) Economic and Social Research Council (ESRC)-funded Doctoral Training Partnership.

## References

1. Allison, T., McCarthy, G., & Wood, C. C. (1992). The relationship between human long-latency somatosensory evoked potentials recorded from the cortical surface and from the scalp. Electroencephalography and Clinical Neurophysiology, 84(4), 301–314. 10.1016/0168-5597(92)90082-m

2. Allison, T., Mccarthy, G., Wood, C. C., & Jones, S. J. (1991). Potentials evoked in human and monkey cerebral cortex by stimulation of the median nerve. A review of scalp and intracranial recordings. Brain, 114, 2465–2503. https://academic.oup.com/brain/article/114/6/2465/371408

3. Alouit, A., Gavaret, M., Ramdani, C., Lindberg, P. G., & Dupin, L. (2025). Cortical activations induced by electrical versus vibrotactile finger stimulation using EEG. NeuroImage, 314. 10.1016/j.neuroimage.2025.121249

4. Auksztulewicz, R., Spitzer, B., & Blankenburg, F. (2012). Recurrent neural processing and somatosensory awareness. Journal of Neuroscience, 32(3), 799–805. 10.1523/JNEUROSCI.3974-11.2012

5. Braun, C., Hess, H., Burkhardt, M., W hle, A., & Preissl, H. (2005). The right hand knows what the left hand is feeling. Experimental Brain Research, 162(3), 366–373. 10.1007/s00221-004-2187-4

6. Cardini, F., Longo, M. R., & Haggard, P. (2011). Vision of the body modulates somatosensory intracortical inhibition. Cerebral Cortex, 21(9), 2014–2022. 10.1093/cercor/bhq267

7. Cauller, L. J., & Kulics, A. T. (1991). The neural basis of the behaviorally relevant N1 component of the somatosensory-evoked potential in SI cortex of awake monkeys: evidence that backward cortical projections signal conscious touch sensation. Experimental Brain Research, 84(3), 607– 619. 10.1007/BF00230973

8. Cohen, L. G., Bandinelli, S., Sato, S., Kufta, C., & Hallett, M. (1991). Attenuation in detection of somatosensory stimuli by transcranial magnetic stimulation. Electroencephalography and Clinical Neurophysiology/Evoked Potentials Section, 81(5), 366–376. 10.1016/0168-5597(91)90026-T

9. Desmedt, J. E., & Tomberg, C. (1989). Mapping early somatosensory evoked potentials in selective attention: critical evaluation of control conditions used for titrating by difference the cognitive P30, P40, P100 and N140. Electroencephalography and Clinical Neurophysiology/Evoked Potentials Section, 74(5), 321–346. 10.1016/0168-5597(89)90001-4

10. Faul, F., Erdfelder, E., Lang, A.-G., & Buchner, A. (2007). G*Power 3: A flexible statistical power analysis program for the social, behavioral, and biomedical sciences. Behavior Research Methods, 39(2), 175–191. 10.3758/BF03193146

11. Forschack, N., Nierhaus, T., Müller, M. M., & Villringer, A. (2020). Dissociable neural correlates of stimulation intensity and detection in somatosensation. NeuroImage, 217. 10.1016/j.neuroimage.2020.116908

12. Gwynne, L., & Tamè, L. (2025). Pain and touch differentially modulate corticospinal excitability, independent of afferent inhibition. 10.1101/2025.05.09.653032

13. Hamada, Y., Otsuka, S., Okamoto, T., & Suzuki, R. (2002). The profile of the recovery cycle in human primary and secondary somatosensory cortex: A magnetoencephalography study. Clinical Neurophysiology, 113(11), 1787–1793. 10.1016/S1388-2457(02)00258-4

14. Hannula, H., Neuvonen, T., Savolainen, P., Tukiainen, T., Salonen, O., Carlson, S., & Pertovaara, A. (2008). Navigated transcranial magnetic stimulation of the primary somatosensory cortex impairs perceptual processing of tactile temporal discrimination. Neuroscience Letters, 437(2), 144–147. 10.1016/j.neulet.2008.03.093

15. Hannula, H., Ylioja, S., Pertovaara, A., Korvenoja, A., Ruohonen, J., Ilmoniemi, R. J., & Carlson, S. (2005). Somatotopic blocking of sensation with navigated transcranial magnetic stimulation of the primary somatosensory cortex. Human Brain Mapping, 26(2), 100–109. 10.1002/hbm.20142

16. Harris, J. A., Harris, I. M., & Diamond, M. E. (2001). The Topography of Tactile Working Memory.

17. Harris, J. A., Miniussi, C., Harris, I. M., & Diamond, M. E. (2002). Transient Storage of a Tactile Memory Trace in Primary Somatosensory Cortex. Journal of Neuroscience, 22(19), 8720–8725.

18. Hirvonen, J., & Palva, S. (2016). Cortical localization of phase and amplitude dynamics predicting access to somatosensory awareness. Human Brain Mapping, 37(1), 311–326. 10.1002/hbm.23033

19. Holmes, N. P., & Tamè, L. (2019). Locating primary somatosensory cortex in human brain stimulation studies: systematic review and meta-analytic evidence. REVIEW Sensory Processing J Neuro-Physiol, 121, 152–162. 10.1152/jn.00614.2018.-Transcranial

20. Holmes, N. P., Tamè, L., Beeching, P., Medford, M., Rakova, M., Stuart, A., & Zeni, S. (2019). Locating primary somatosensory cortex in human brain stimulation studies: experimental evidence. J Neurophysiol, 121, 336–344. 10.1152/jn.00641.2018.-Transcranial

21. Iwamura, Y., Tanaka, M., Iriki, A., Taoka, M., & Toda, T. (2002). Processing of tactile and kinesthetic signals from bilateral sides of the body in the postcentral gyrus of awake monkeys. Behavioural Brain Research, 135(1–2), 185–190. www.elsevier.com/locate/bbr

22. Jones, S. R., Pritchett, D. L., Stufflebeam, S. M., Hämäläinen, M., & Moore, C. I. (2007). Neural correlates of tactile detection: A combined magnetoencephalography and biophysically based computational modeling study. Journal of Neuroscience, 27(40), 10751–10764. 10.1523/JNEUROSCI.0482-07.2007

23. Katus, T., Müller, M. M., & Eimer, M. (2015). Sustained Maintenance of Somatotopic Information in Brain Regions Recruited by Tactile Working Memory. The Journal of Neuroscience, 35(4), 1390– 1395. 10.1523/JNEUROSCI.3535-14.2015

24. Koch, G., Franca, M., Albrecht, U. V., Caltagirone, C., & Rothwell, J. C. (2006). Effects of paired pulse TMS of primary somatosensory cortex on perception of a peripheral electrical stimulus. Experimental Brain Research, 172(3), 416–424. 10.1007/s00221-006-0359-0

25. Kulics, A. T. (1982). Cortical neural evoked correlates of somatosensory stimulus detection in the rhesus monkey. Electroencephalography and Clinical Neurophysiology, 53(1), 78–93. 10.1016/0013-4694(82)90108-0

26. Kulics, A. T., & Cauller, L. J. (1986). Cerebral cortical somatosensory evoked responses, multiple unit activity and current source-densities: their interrelationships and significance to somatic sensation as revealed by stimulation of the awake monkey’s hand. Experimental Brain Research, 62(1). 10.1007/BF00237402

27. Lamotte, R. H., & Mountcastle, V. B. (1979). Disorders in Somethesis Following Lesions of Parietal Lobe. Neurophysiology, 42(2), 400–419.

28. Levitt, H. (1971). Transformed up-down methods in psychoacoustics. The Journal of the Acoustical Society of America, 49(2B), 467–477.

29. Libet, B., Alberts, W. W., Wright, E. W., & Feinstein, B. (1967). Responses of Human Somatosensory Cortex to Stimuli below Threshold for Conscious Sensation. Science, 158(3808), 1597–1600. 10.1126/science.158.3808.1597

30. Mauguière, F., Merlet, I., Forss, N., Vanni, S., Jousmäki, V., Adeleine, P., & Hari, R. (1997). Activation of a distributed somatosensory cortical network in the human brain: a dipole modelling study of magnetic fields evoked by median nerve stimulation. Part II: Effects of stimulus rate, attention and stimulus detection. Electroencephalography and Clinical Neurophysiology, 104(4), 290–295. 10.1016/s0013-4694(97)00018-7

31. McKay, D. R., Ridding, M. C., & Miles, T. S. (2003). Magnetic stimulation of motor and somatosensory cortices suppresses perception of ulnar nerve stimuli. International Journal of Psychophysiology, 48(1), 25–33. 10.1016/S0167-8760(02)00159-9

32. Morley, J. W., Vickery, R. M., Stuart, M., & Turman, A. B. (2007). Suppression of vibrotactile discrimination by transcranial magnetic stimulation of primary somatosensory cortex. European Journal of Neuroscience, 26(4), 1007–1010. 10.1111/j.1460-9568.2007.05729.x

33. Mountcastle, V. B., Talbot, W. H., Sakata, H., & Hyvärinen, J. (1969). Cortical Neuronal Mechanisms in Flutter-Vibration Studied in Unanesthetized Monkeys. Neuronal Periodicity and Frequency Discrimination. Journal of Neurophysiology, 32(3), 452–484.

34. Palva, S., Linkenkaer-Hansen, K., Näätänen, R., & Palva, J. M. (2005). Early neural correlates of conscious somatosensory perception. Journal of Neuroscience, 25(21), 5248–5258. 10.1523/JNEUROSCI.0141-05.2005

35. Peterson, N. N., Schroeder, C. E., & Arezzo, J. C. (1995). Neural generators of early cortical somatosensory evoked potentials in the awake monkey. In Electroencephalography and clinical Neurophysiology (Vol. 96).

36. Romo, R., Hernández, A., Zainos, A., Brody, C. D., & Lemus, L. (2000). Sensing without Touching. Neuron, 26(1), 273–278. 10.1016/S0896-6273(00)81156-3

37. Rossi, S., Antal, A., Bestmann, S., Bikson, M., Brewer, C., Brockmöller, J., Carpenter, L. L., Cincotta, M., Chen, R., Daskalakis, J. D., Di Lazzaro, V., Fox, M. D., George, M. S., Gilbert, D., Kimiskidis, V. K., Koch, G., Ilmoniemi, R. J., Pascal Lefaucheur, J., Leocani, L., … Hallett, M. (2021). Safety and recommendations for TMS use in healthy subjects and patient populations, with updates on training, ethical and regulatory issues: Expert Guidelines. Clinical Neurophysiology, 132(1), 269–306. 10.1016/j.clinph.2020.10.003

38. Rossini, P. M., Barker, A. T., Berardelli, A., Caramia, M. D., Caruso, G., Cracco, R. Q., Dimitrijević, M. R., Hallett, M., Katayama, Y., Lücking, C. H., Maertens de Noordhout, A. L., Marsden, C. D., Murray, N. M. F., Rothwell, J. C., Swash, M., & Tomberg, C. (1994). Non-invasive electrical and magnetic stimulation of the brain, spinal cord and roots: basic principles and procedures for routine clinical application. Report of an IFCN committee. Electroencephalography and Clinical Neurophysiology, 91(2), 79–92. 10.1016/0013-4694(94)90029-9

39. Schröder, P., Nierhaus, T., & Blankenburg, F. (2021). Dissociating perceptual awareness and postperceptual processing: The P300 is not a reliable marker of somatosensory target detection. Journal of Neuroscience, 41(21), 4686–4696. 10.1523/JNEUROSCI.2950-20.2021

40. Schubert, R., Blankenburg, F., Lemm, S., Villringer, A., & Curio, G. (2006). Now you feel it – Now you don’t: ERP correlates of somatosensory awareness. Psychophysiology, 43(1), 31–40. 10.1111/j.1469-8986.2006.00379.x

41. Seyal, M., Masuoka, L. K., & Browne, J. K. (1992). Suppression of cutaneous perception by magnetic pulse stimulation of the human brain. Electroencephalography and Clinical Neurophysiology/Evoked Potentials Section, 85(6), 397–401. 10.1016/0168-5597(92)90053-E

42. Seyal, M., Siddiqui, I., & Hundal, N. S. (1997). Suppression of spatial localization of a cutaneous stimulus following transcranial magnetic pulse stimulation of the sensorimotor cortex. Electroencephalography and Clinical Neurophysiology/Electromyography and Motor Control, 105(1), 24–28. 10.1016/S0924-980X(96)96090-7

43. Soininen, K., & Järvilehto, T. (1983). Somatosensory evoked potentials associated with tactile stimulation at detection threshold in man. Electroencephalography and Clinical Neurophysiology, 56(5), 494–500. 10.1016/0013-4694(83)90234-1

44. Tamburin, S., Manganotti, P., Zanette, G., & Fiaschi, A. (2001). Cutaneomotor integration in human hand motor areas: Somatotopic effect and interaction of afferents. Experimental Brain Research, 141(2), 232–241. 10.1007/s002210100859

45. Tamè, L., Carr, A., & Longo, M. R. (2017). Vision of the body improves inter-hemispheric integration of tactile-motor responses. Acta Psychologica, 175, 21–27. 10.1016/j.actpsy.2017.02.007

46. Tamè, L., & Holmes, N. P. (2016). Involvement of human primary somatosensory cortex in vibrotactile detection depends on task demand. NeuroImage, 138, 184–196. 10.1016/j.neuroimage.2016.05.056

47. Tamè, L., & Holmes, N. P. (2023). Neurostimulation in Tactile Perception. In Somatosensory Research Methods (pp. 451–482). 10.1007/978-1-0716-3068-6_20

48. Tamè, L., Pavani, F., Braun, C., Salemme, R., Farnè, A., & Reilly, K. T. (2015). Somatotopy and temporal dynamics of sensorimotor interactions: Evidence from double afferent inhibition. European Journal of Neuroscience, 41(11), 1459–1465. 10.1111/ejn.12890

49. Tamè, L., Pavani, F., Papadelis, C., Farnè, A., & Braun, C. (2015). Early integration of bilateral touch in the primary somatosensory cortex. Human Brain Mapping, 36(4), 1506–1523. 10.1002/hbm.22719

50. Tsujinaka, R., Oda, H., Fukuda, S., Hamada, N., Matsuoka, M., & Hiraoka, K. (2023). Afferent volley from the digital nerve induces short-latency facilitation of perceptual sensitivity and primary sensory cortex excitability. Experimental Brain Research, 241(5), 1339–1351. 10.1007/s00221-023-06611-y

51. Wühle, A., Mertiens, L., Rüter, J., Ostwald, D., & Braun, C. (2010). Cortical processing of near-threshold tactile stimuli: An MEG study. Psychophysiology, 47(3), 523–534. 10.1111/j.1469-8986.2010.00964.x

52. Wühle, A., Preissl, H., & Braun, C. (2011). Cortical processing of near-threshold tactile stimuli in a paired-stimulus paradigm – an MEG study. European Journal of Neuroscience, 34(4), 641–651. 10.1111/j.1460-9568.2011.07770.x

