## Supplemental Figure S1 for "Temporal constraints of conscious tactile perception in the primary somatosensory cortex"

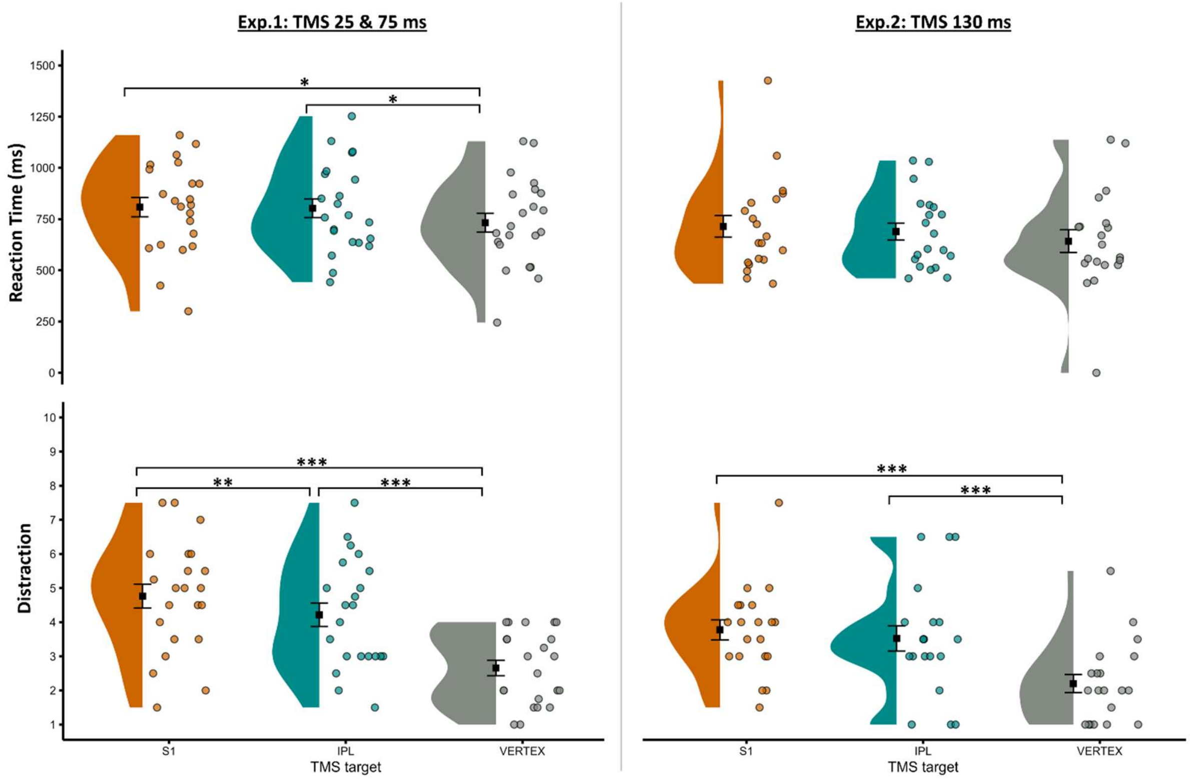


**Figure S1.** Tactile detection reaction times and TMS distraction ratings as a function of TMS target when early TMS was delivered after afferent onset (25 & 75 ms, Experiment 1) or, late TMS delivery (130 ms, Experiment 2). Reaction times are given for "hits" (a "go" response on target-present trials). Black squares show mean and standard error means (SEM). Half violins show data distribution, coloured circles are jittered individual participant data points*$P \leq .05$, **$P \leq .01$, ***$P \leq .001$
